# A tale of two phage tails: Engineering the host range of bacteriophages infecting *Clostridioides difficile*

**DOI:** 10.1101/2023.10.16.562632

**Authors:** Joanna P. Steczynska, Sarah J. Kerr, Michelle L. Kelly, Michaella J. Whittle, Terry W. Bilverstone, Nigel P. Minton

## Abstract

*Clostridioides difficile* infection (CDI) is a leading cause of antibiotic-associated diarrhoea across the globe. Although treatable with a restricted number of antibiotics, the emergence of resistant variants and high relapse rates necessitate alternative countermeasures. Phage therapy represents an attractive option. However, its implementation is handicapped by the narrow host specificity of the *C. difficile* bacteriophages isolated to date. One strategy to rationally expand phage host range would be to make appropriate modifications to the phage receptor binding protein (RBP). Here, we identify the tail fibre as the RBP of two *Myoviridae* phages, ΦCD1801 and ΦCD2301, which were previously isolated and propagated using the *C. difficile* strains CD1801 (RT078) and CD2301 (RT014), respectively. Contrary to studies into reprogramming the host ranges of phage of other bacterial other species, exchanging the tail fibre genes (*tcf/tfp*) alone between the two phage was insufficient to change host specificity. Rather, alterations to host range were dependent their exchange together with a putative chaperone encoded by *hyp*, localised adjacent to the tail fibre gene. Capitalising on this discovery, CRISPR/Cas9 was used to change the host range of one phage to that of the other by swapping the respective *tcf/tfp* and *hyp* genes. Significantly, one of the resulting mutants, surpassed both parental phages in terms of host range and efficiency of infection. This is the first time that genome engineering has successfully expanded the host range of a *C. difficile* phage, a prerequisite for implementing phage for the treatment of CDI.

**Importance:** Alternatives to antibiotics for treating *Clostridioides difficile* infection (CDI) are urgently required. Phage therapy presents an attractive option as it has the potential to clear the infection with minimal microbiome disruption and eliminate the possibility of recurrence. However, the *C. difficile* bacteriophages isolated to date have highly restricted host ranges. Moreover, rational strategies to alter specificity have till now been precluded as the identity of the phage receptor binding proteins involved was largely unknown. Here, we demonstrated that tail fibre proteins and an associated putative chaperone determine the host range of two *Myoviridae* phage. This enabled the alteration of specificity through CRISPR-mediated genome editing and the creation of a phage derivative with a host range and infection efficiency exceeding that of the parental phages. This is the first time that the host range of a *C. difficile* phage has been successfully expanded through rational genome engineering.

## Introduction

The pathogenic anaerobe, *Clostridioides difficile* (formerly *Clostridium difficile*), is considered a major threat to public health by the US Centre for Disease control and Prevention which demands urgent attention ^1^. The rising case rates of *Clostridioides difficile* infections (CDI) over the last 20 years have led to it becoming the most common hospital-associated infection ^2^. In the UK alone, ∼43,000 cases were reported for the 2021/22 financial year. CDI occurs following prolonged antibiotic treatment which disrupts the integrity of the gut microbiome, leading to dysbiosis and proliferation of the pathogen^3^. The complications that arise can sometimes be fatal^4^. Further, the spore-forming nature of *C. difficile* leads to relapse in up to 35% of patients^5^ with some individuals suffering multiple recurrences. Paradoxically, the first-line therapy for CDI is antibiotic treatment^6^. The continued long-term use of antibiotic therapy may lead to reduced susceptibility of the bacterium to antimicrobials and further instigate emergence of resistant and hypervirulent strains^7,8^. The nature of CDI calls for a more targeted treatment with precise therapies that leave the microbiome unperturbed.

A particularly attractive option would be the implementation of phage therapy. This is because bacteriophages (phages), the most abundant entities on Earth^9^, are known for their precise host specificity. This can range from the infection of an entire genus, down to subspecies level^10^. Phages have been used for over 100 years to cure infections, although largely superseded in the West after 1940 with the development of antibiotics^11,12^. Phages therefore represent a promising, narrow-spectrum alternative to antibiotics for the treatment of CDI – clearing the infection with minimal microbiome disruption and eliminating the possibility of recurrence. Isolated phages of *C. difficile*, however, are generally known for their very narrow host ranges, although some “broad range” phages infecting prevalent ribotypes have been isolated^13,14^. Their specificity is a welcome advantage, but also a double-edged sword as the currently available phages are not able to target all clinically prevalent *C. difficile* strains. To address the issue of specificity, the underlying mechanisms dictating phage binding range require investigation. Phage receptor binding protein (RBP) and host range determining factors are poorly characterised in phages infecting *C. difficile*, although it has been established that the counterpart host receptor is encoded by the *slpA* gene^14–18^. Part of the phage host range is its binding range, dictated by the tail components, mainly the RBP^19–22^, which is recognised as the tail fibre (or spike in other species)^23^. Many phages also require accessory proteins to support tail fibre function ^24–27^, although genes of this nature have not yet been reported in *C. difficile* phages.

To overcome the current barriers to phage therapy in *C. difficile*, we sought to establish the receptor binding determinants of the typical *C. difficile Myoviridae* phages and thereafter alter host range through genome engineering. This was accomplished through the use of a CRISPR/Cas9-based editing tool, RiboCas^29^, to create seamless deletion, complement and exchange mutants. The latter comprised phage mutants in which their tail genes were replaced with those of a second phage with different host specificity. Testing of the ability of the hybrid phages to infect a range of *C. difficile* strains confirmed the tail fibre as the principal determinant of host specificity and revealed supporting tail genes. This study provides a blueprint for host range manipulation of *Myoviridae* phages infecting *C. difficile*.

## Results

### Tail fibre validation

Our strategy was based on two phages, ΦCD1801 and ΦCD2301, which were previously isolated and propagated using the *C. difficile* strains CD1801 (RT078) and CD2301 (RT014), respectively ^14^. The initial focus was on the phage tail fibre-encoding genes, before expanding to include other tail genes within the vicinity. First, we sought to establish the identity of the genes encoding the RBP, usually the tail fibre, and thereafter determine whether they dictate binding specificity.

Tail fibre genes are typically located between the genes encoding the tail tape measure protein and the endolysin^18,30,31^ and fall within the structural tail gene module^32^.A BLAST query of genes located within the tail module revealed the presence of “tail collar fiber protein” (QVW56628.1, *tcf*) in ΦCD1801 and “tail protein” (QVW56667.1, hereafter referred to as *tail fibre protein* – *tfp*) in ΦCD2301; both ∼800bp in size. These were predicted to be the RBP for each phage through homology and location. The two proteins shared 83% identity and 92% similarity, exhibiting the greatest divergence at their C-terminals (Supplementary, S5). Using HMMER^33^, both were shown to contain a “domain of unknown function” (DUF3751), at the N-terminal (Accession IPR022225). This domain is a common feature of phage tail fibres across many species^33,34^ and docks the fibre to the baseplate^35^. The C-terminal region likely determines the binding specificity and is, therefore, less conserved between phages. The same pattern can be found between this region of ΦCD1801 and ΦCD2301, which likely explains their differing host ranges^14^.

To enable functional verification of the identified tail fibre genes, RiboCas was used to generate deletion mutants of both phages (Fig. 1a). The resultant two mutant phages lacking their respective tail fibre genes (ΦCD1801ΔBM1 and ΦCD2301ΔBM1), were then assessed (Fig. 1b). No binding was detected in either case as no plaques were obtained. Binding capabilities were restored to that of the WT phages (Fig. 2a and 2b) following complementation of the deleted gene in each phage (ΦCD1801*CompBM1 and ΦCD2301*CompBM1).

**Fig. 1.**
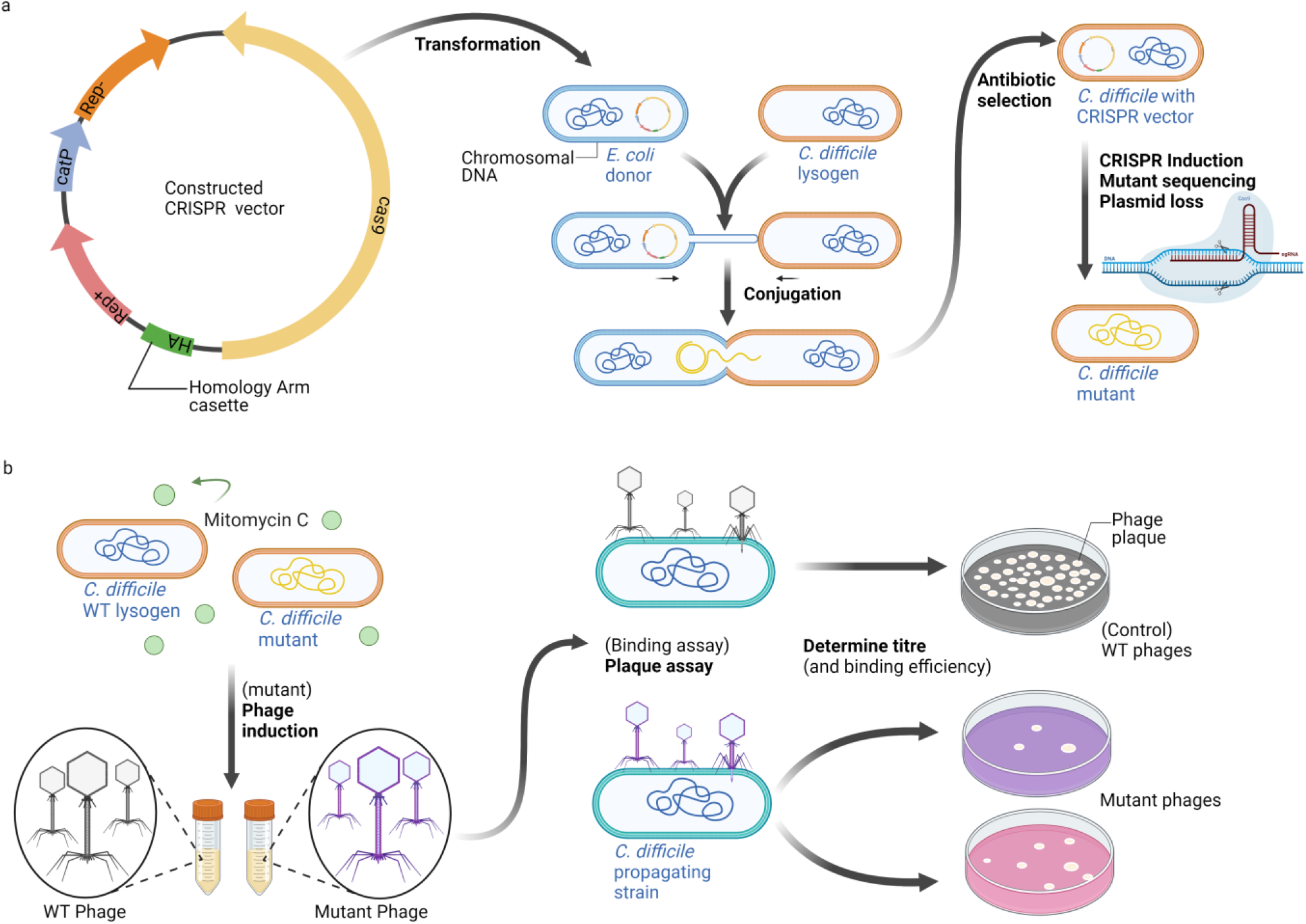
Mutant generation and evaluation. **a)** CRISPR/Cas9 based editing using RiboCas. The constructed vector (left) was transformed into an *E. coli* donor and conjugated, then screened for the mutation (right). **b)** Mutant testing. The mutant was then induced to produced phages (left), which were enumerated via plaque assay (right). The plaques were counted, and the titre was determined for each mutant, alongside WT controls. As appropriate, a binding assay was carried out alongside a standard plaque assay to determine degree of binding. Figure created using BioRender.com.

**Fig. 2.**
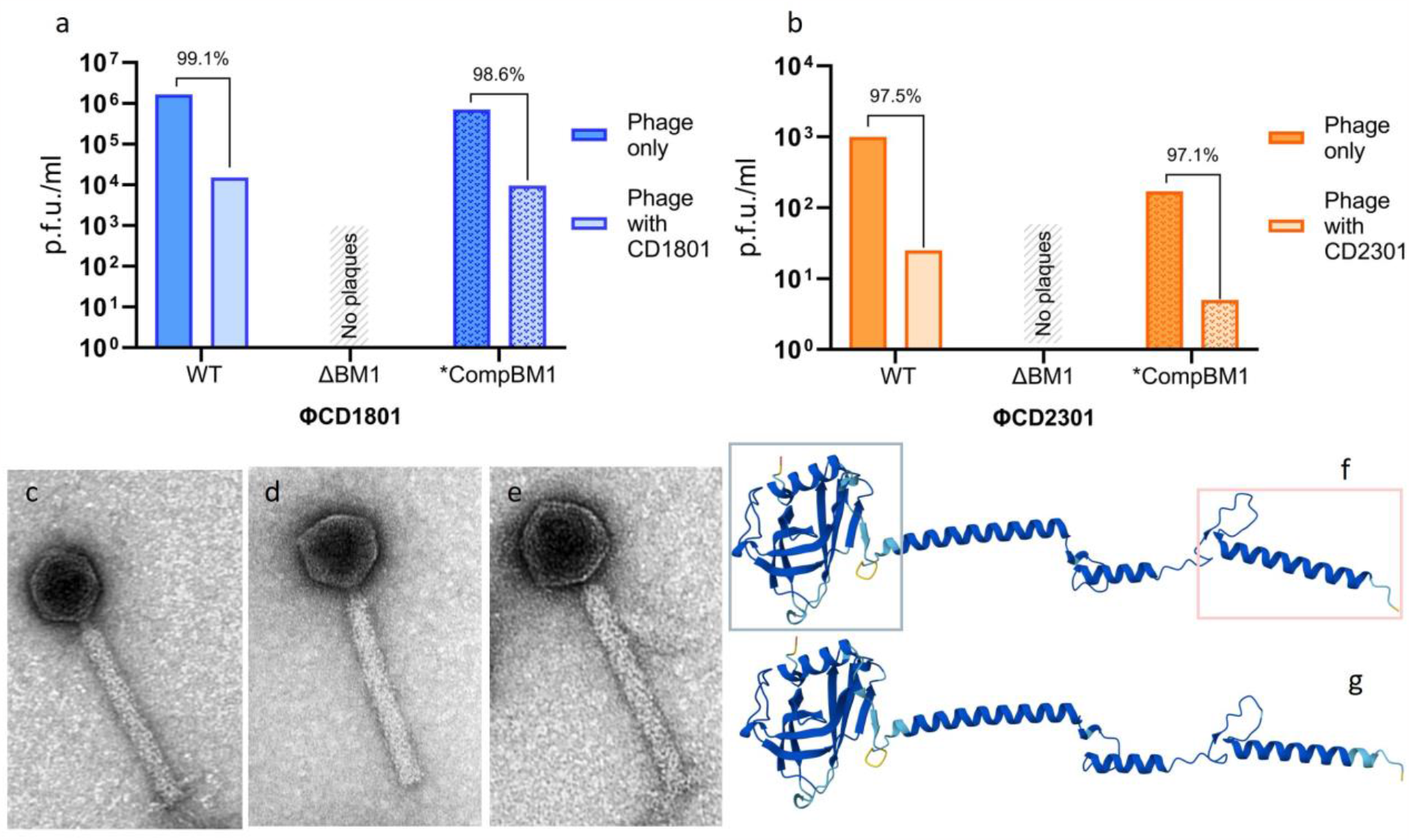
Validation of tail fibres. **a & b)** Illustrated are the relative degree of binding of WT and mutant phages to their respective hosts. The WT and Complement (ΦCD1801*CompBM1 and ΦCD2301*CompBM1) mutants show a high degree of binding (>97%), whereas plaquing, and thus binding, was not detected in the ΔBM1 mutants. Panels c-e show the morphology of the phage particles obtained using TEM. **c)** TEM of the WT ΦCD1801 phage with tail fibre-like projections at the terminal end of the tail. **d)** TEM of a ΦCD1801ΔBM1 phage particle, where the tail fibres have been deleted, represented by the smooth, blunt terminal-tail appearance. Images presented are representative of the morphology observed for each sample. **e)** TEM of the ΦCD1801*CompBM1 complemented mutant. **f & g)** Shows a AlphaFold 2 prediction of the tertiary structures of the respective tail fibres (1f-*tcf* in ΦCD1801; 1g-*tfp* in ΦCD2301). The structures exhibit a relatively high confidence score (pLDDT > 90, dark blue) for majority of the prediction. Areas in light blue and yellow are of lower confidence (90 > pLDDT > 50) scores, although not prevalent. The structure shows the DUF3751 domain, as well as the long fibre-like projection, created by the α-helices.

The examination of mutant and WT phage particles by TEM showed that ΦCD1801ΔBM1 was able to form complete virion particles that maintain structural integrity. When compared to the WT and ΦCD1801*CompBM1 phage particles, however, the deletion mutant appeared to lack tail fibres (Fig. 2c-e). This was indicated by the “blunt” appearance of ΦCD1801ΔBM1 tail ends (Fig. 2d), whereas the presence of fibres in WT and ΦCD1801*CompBM1 phage particles gave their tails a “fluffy” appearance (Fig. 2c and 2e). Further, using AlphaFold 2^36^, the 3D protein structure of the tail fibres was predicted (Fig. 2f and 2g). The resulting predictions resemble a tail fibre comprised of the DUF3751 domain (attaches fibre to the baseplate) mostly depicted by tightly packed β-sheets, which leads onto α-helices arranged into a long fibre-like projection (that recognises and binds the receptor). Combined, these data strongly suggest that these genes encode tail fibres, recognised as the phage RBP.

### Mapping of phage genes involved in binding

As binding range is determined by the tail fibre and often supported by neighbouring tail proteins, we sought to manipulate it by generating hybrid mutants of ΦCD1801 in which various tail genes (termed ‘binding modules’) were exchanged with the equivalent genes from ΦCD2301 and vice versa. Although the overall sequence identity between the ΦCD1801 and ΦCD2301 genomes is low, at only 36%, the tail regions share 71.6% identity and appear highly homologous. Despite the latter homology, the two phages do not have a shared host range. First, we mapped the tail region of both phages to identify additional genes, apart from the tail fibre (encoded by *tcf* and *tfp*), that may play a direct or supporting role in the host range encoding the following components: the baseplate (*bpJ*), tail component (*ptp* and *ptc*) and hypothetical protein (*hyp*). Single deletion and exchange mutants were generated to help resolve the individual contribution of each gene. We anticipated that the exchange of multiple genes may be required to maintain structural integrity (upstream genes – *bpJ*; *ptp* and *ptc*) and/or provide an accessory function like a chaperone (downstream gene – *hyp*). In total, seven regions/genes were targeted for testing – designated as Binding Module (BM) 1 to 7 (Fig. 3).

**Fig. 3.**
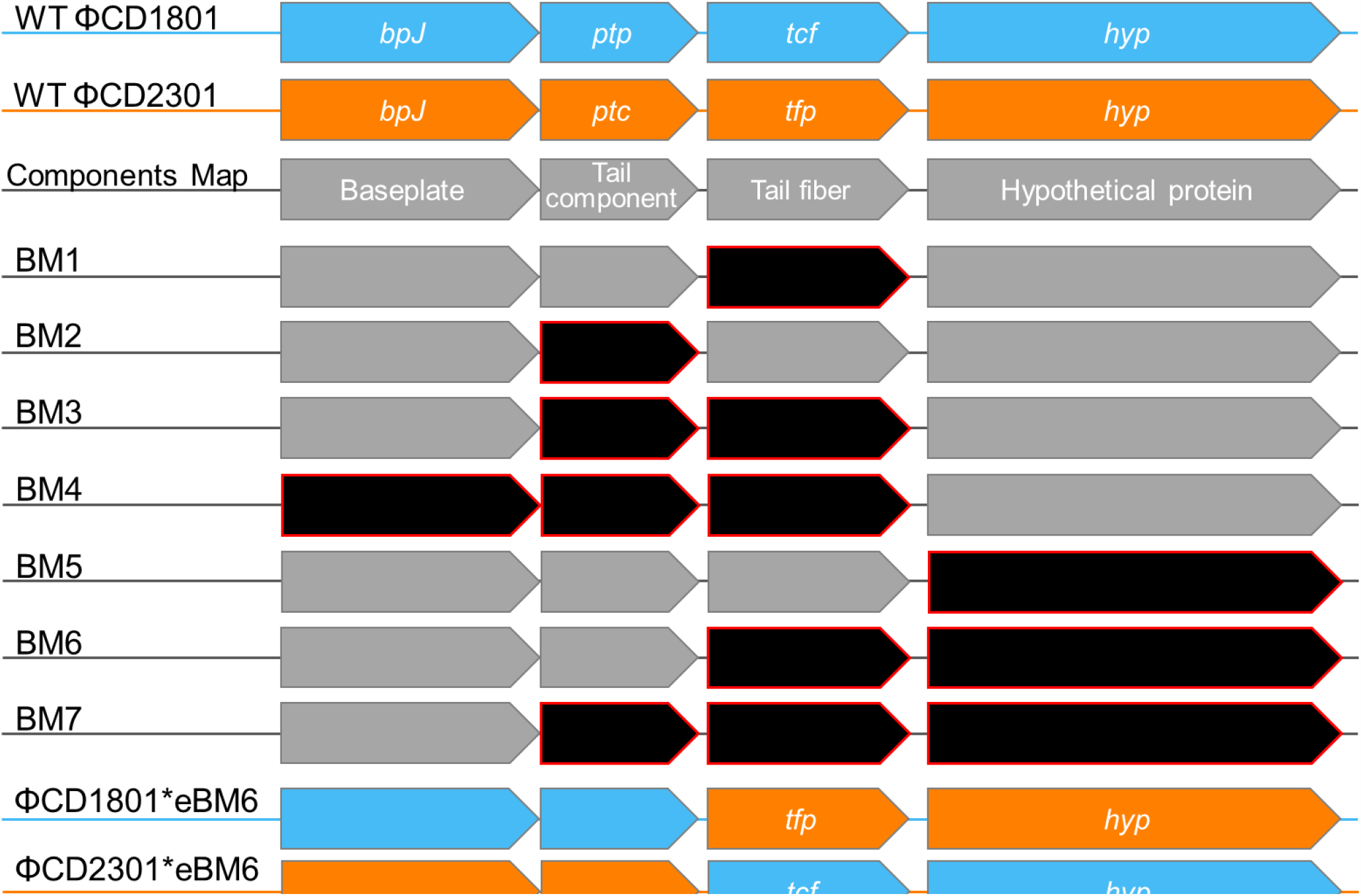
Map of the tail gene locus. The components map represents the four genes that were investigated in this study. The full gene names are as follows: *bpJ =* baseplate protein J, *ptp =* putative tail protein, *ptc =* putative tail component, *tcf =* tail collar fibre protei*n, tfp =* tail (fibre) protei*n, hyp* = hypothetical protein. *The ‘tail protein’ gene of ΦCD2301 was renamed as ‘tail fibre protein’, following its confirmation as such in this study. Illustrated are the genes affected in each deletion and exchange (BM1-7) mutant in black with red borders; where relevant genes and backbone encoded by ΦCD1801 are colour-coded in light blue, and those encoded by ΦCD2301 in orange. Genes coloured in grey were unaffected. The gene fragments are representative of the size and location of the actual genes.

BM1, investigated in the preceding experiments, comprised the tail fibre gene and was expected to be the main determinant of host range. BM2-4 mutants were generated to ensure structural compatibility between the tail fibre and upstream components like the tail component (BM2 and BM3) and baseplate (BM4). Many phages utilise chaperones to fold tail fibres ^37–39^, and in some cases, ensure their function ^24,27^. Consequently, BM5 and 6 mutants were created to determine whether the hypothetical protein could play a role in tail fibre function. BM7 (composed of the tail component, tail fibre and the hypothetical protein) was included in case both structural and chaperone activities were at play.

### Identification of binding modules that determine host range through deletions and exchanges

We generated further deletion (ΦCD1801ΔBM2-7 and ΦC23801ΔBM2-7), and subsequently, exchange mutants (ΦCD1801*eBM1-6 and ΦC23801*eBM1-7) to determine the combination of genes needed to change phage host range.

Out of all 14 deletion mutants generated, only mutant ΦCD1801ΔBM5 was able to bind to its original propagating host – CD1801 (Fig. 4a). The phage’s binding capability, however, was substantially diminished when compared to the WT, showing a 99.98% decrease in number of particles which were able to bind. This effect was not observed in the parallel mutant – ΦCD2301ΔBM5 (Fig. 4b) and is likely a consequence of a much lower induction titre of ΦCD2301 at 10^4^ compared to 10^7^ pfu/ml of ΦCD1801. The hypothetical titre of ΦCD2301ΔBM5 would therefore be expected to be reduced to 3.5 pfu/ml, which is below the experimental detection limit (10 pfu/ml).

**Fig. 4.**
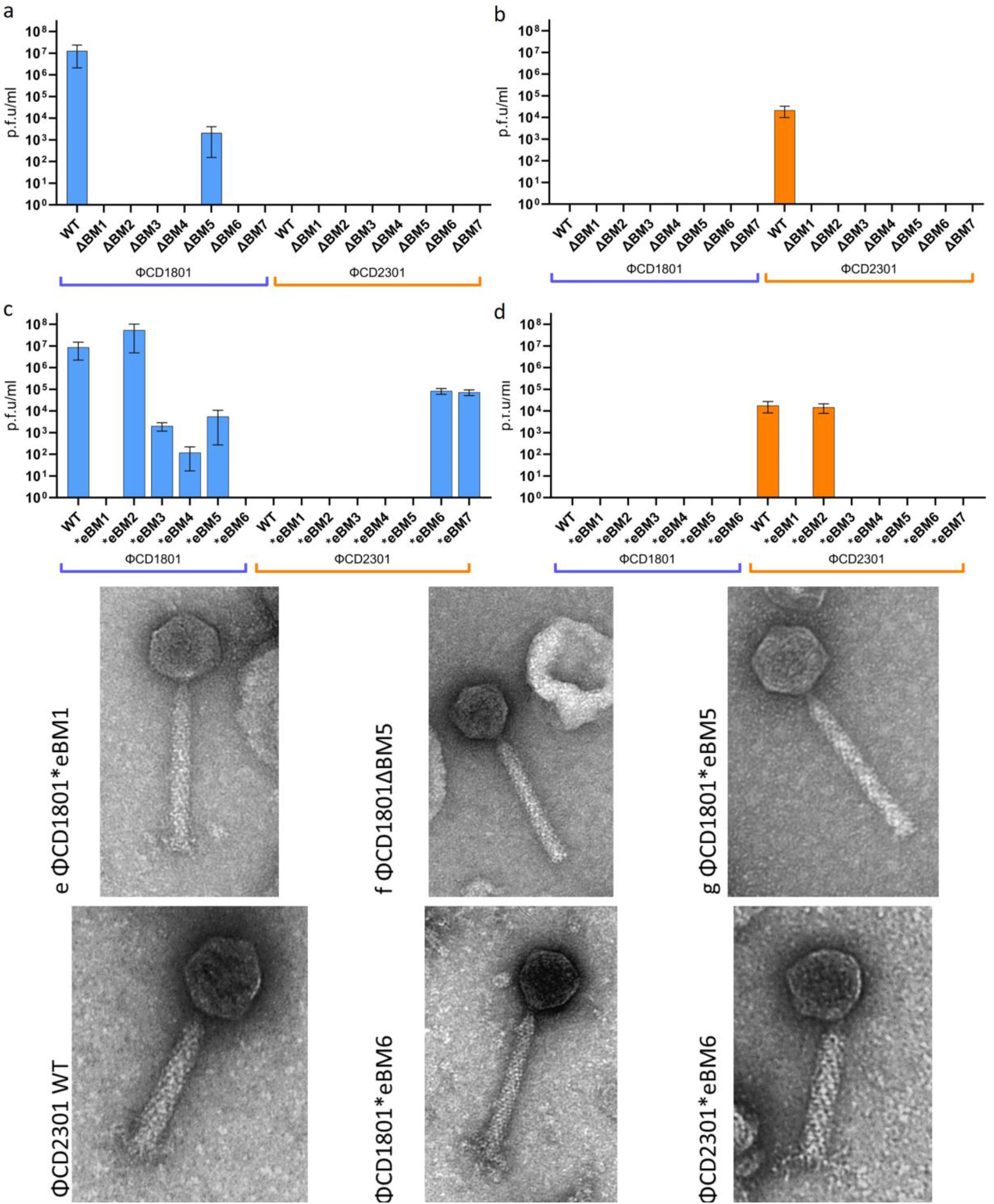
Plaquing and TEM imaging of mutants. **a & b)** the pfu/ml of the deletion mutants plaqued against CD1801 and CD2301 bacterial hosts, respectively. **c & d)** the pfu/ml of the exchange mutants plaqued against CD1801 and CD2301 bacterial hosts, respectively. **e-j)** morphology of phage particles obtained using TEM. The tails in panels f and g appear blunt, compared to the fluffy tails seen in h-j. The tail in panel e appears less fluffy at the bottom compared to the WT phage (2c) but seems to have some sort of tail structure.

Following, exchange mutants were generated by inserting the binding module genes from one phage into the other in place of the previously deleted genes. The backbone of the exchange mutants remained unchanged and the resulting hybrid phage mutants (ΦCD1801*eBM1-6 and ΦCD2301*eBM1-7) now carried various binding modules from the other phage (Fig. 3). Generation of the ΦCD1801*eBM7 mutant proved not to be possible, despite multiple attempts, likely owing to challenges in the uptake of large vectors by CD1801. The mutant exchange phages generated were then induced and the ability of the released phage particles to infect either propagating strain (CD1801 and CD2301) was tested by plaque assay (Fig. 1b).

Exchange of the tail fibres alone (ΦCD1801*eBM1 and ΦCD2301*eBM1) did not result in infectious phage particles as the mutants were unable to infect either of the propagating hosts (Fig. 4c and 4d). Under TEM, ΦCD1801*eBM1 appeared to maintain structural integrity showing complete virion particles with “fluffy” tails (Fig. 4e), indicating a tail fibre was present, yet lacking in function.

Mutant ΦCD1801*eBM2 and its counterpart ΦCD2301*eBM2 were both able to infect their original propagating strains at levels similar to that of the WT phages (no statistical significance). Combined with the lack of infection when the tail component is deleted (ΔBM2) this result showed that the tail component gene needs to be present for a functional phage particle to be formed, however, it is non-specific and interchangeable between these two phages.

Mutants ΦCD1801*eBM3 and ΦCD1801*eBM4 were also able to plaque, albeit at extremely low levels (99.98% and 99.99% decrease, respectively) and again only on the original propagating strain, CD1801. These data indicate that the structural genes targeted in BM2-4 do not play a role in determining host range as the mutants were unable to plaque on their respective new target strains. Mutant ΦCD1801*eBM5 was able to bind to CD1801 at levels comparable to those of the deletion mutant (ΦCD1801ΔBM5), showing that the hypothetical protein originating from ΦCD2301 is not functional in ΦCD1801 on its own and appears specific to the tail fibre of origin. Further, under TEM the deletion and exchange mutants both appear blunt (Fig. 4f and g), as if the tail fibres are not present or folded correctly. The reciprocal mutants ΦCD2301*eBM3-5 did not bind to either strain. This was expected, however, due to the previously specified low induction level of the phage putting mutants below the detection limit.

Significantly, the exchange mutants ΦCD2301*eBM6 and ΦCD2301*eBM7 were able to plaque on the propagating strain of ΦCD1801, CD1801. This represents the first time that the host range of a *C. difficile* phage has been deliberately changed. Both mutants encoded the tail fibre, as well as the downstream hypothetical protein from ΦCD1801. Mutant ΦCD2301*eBM7 additionally encoded the tail component from ΦCD1801, which we previously showed through *eBM2 to not be involved in specificity. Consequently, its exchange in future experiments is unnecessary. Taken together, these data indicate that the tail fibre and its corresponding downstream hypothetical protein determine binding specificity.

Given the change in host range observed with ΦCD2301*eBM6, the reciprocal exchange mutant ΦCD1801*eBM6 would have been expected to infect CD2301, however this was not the case. As this apparent barrier may be specific to strain CD2301, we tested several other strains previously shown to be infected by WT ΦCD2301^14^. Of those tested, strain CD8001 was found to be particularly susceptible to ΦCD1801*eBM6 and was therefore designated as the new propagating host in following experiments. This indicated a problem with entry of ΦCD1801 backbone into CD2301 cells specifically, rather than the exchanged tail components. The phage particles were imaged under TEM, as the titre of the mutants could be increased to an appropriate level. The WT phages (ΦCD1801 – 2c, ΦCD2301 – 4h) and their respective *eBM6 mutants (Fig. 4i and j) all appeared to have fibres at the terminal tail ends; they could also be differentiated by their backbones as ΦCD1801 tails are ∼160nm in length, but ΦCD2301 tails are much shorter at ∼100nm.

### Exchange mutants mimic host range of WT phages based on tail components

We next tested the host range of the ΦCD1801*eBM6 and ΦCD2301*eBM6 exchange mutants, alongside the WT phages, against a panel of *C. difficile* isolates (Fig. 5). The panel comprised 75 strains selected based on previously infected ribotypes (RT002, 014, 078 and 122) and previous susceptibility to infection by other phages. The titre of each preparation of phage particles in the panel was normalised to 10^8^ pfu/ml using their propagating strain.

**Fig. 5.**
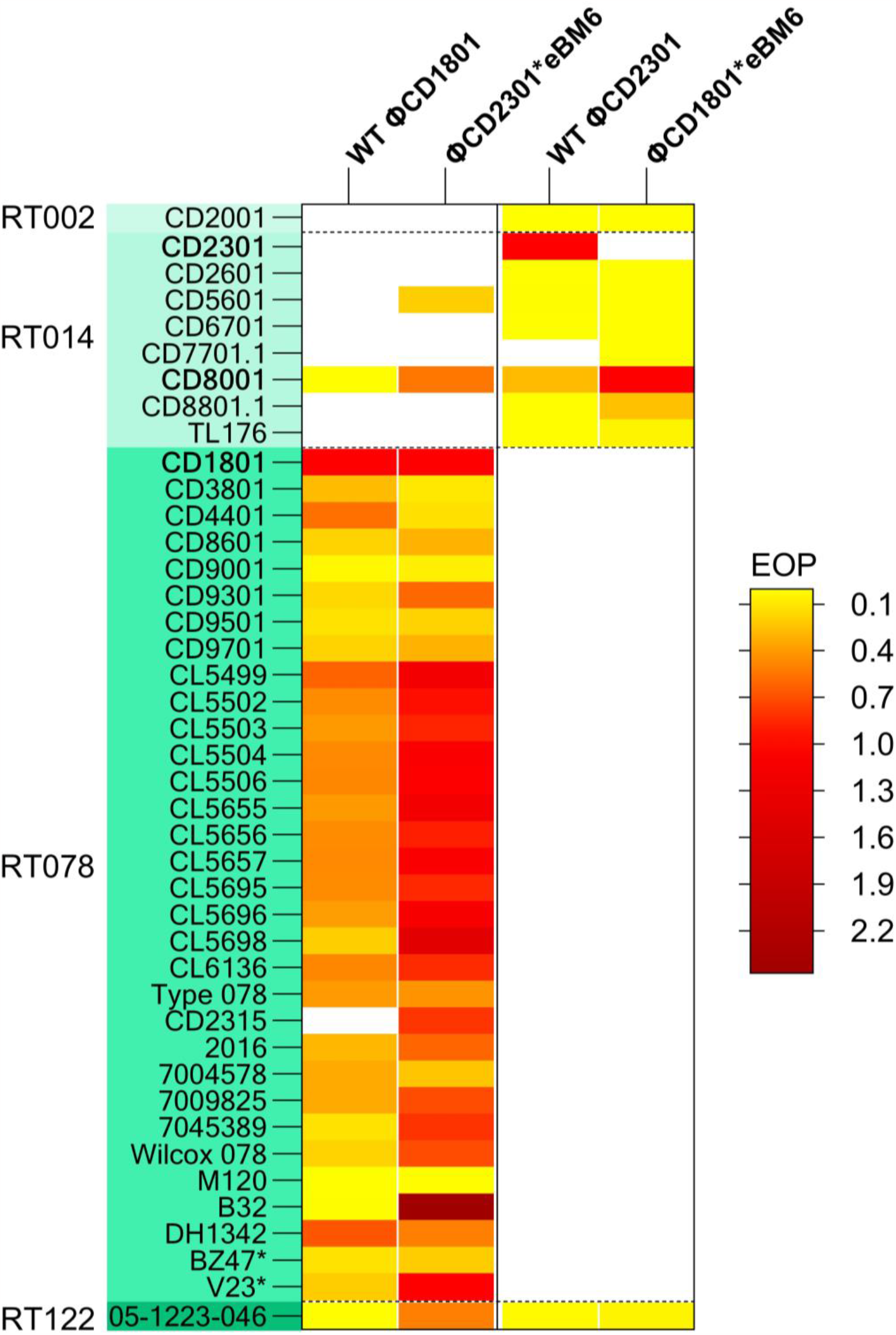
Host range heatmap. The data illustrated is for the WT parent phages as well as their respective *eBM6 mutants. The colour scale is indicative of the degree of infection when compared with the propagating strain, calculated as EOP – efficiency of plating. Yellow coloured rectangles, mostly populated by ΦCD2301 and mutant following its host range, ΦCD1801*eBM6, indicate very inefficient infection. Whereas the red, and deep red coloured rectangles indicate a much more efficient infection. Propagating strains are highlighted in bold. The EOP of propagating strains with corresponding phages is always 1. The full list of strains used, their origins and corresponding EOP values can be found in the Supplementary data. White rectangles indicate no infection was detected. *indicates ribotype 786, see Supplementary, S4.

As expected, mutant ΦCD2301*eBM6 was able to fully mimic the host range of WT ΦCD1801 and infected 35 out of 75 strains, including strains CD7701.1 and CD2315 which were not infected by the WT phage. Strain CD2315 (RT078) was previously shown to exhibit lysogenic immunity to WT ΦCD1801 ^14^, however ΦCD2301*eBM6 was able to infect it at EOP 0.79 (Supplementary, S6) as the repressor found in its ΦCD2301 backbone differs considerably to that of WT ΦCD1801 (19% identity). Moreover, the EOP of the mutant phage on the strains was more than twice that of the WT ΦCD1801, being 0.70 and 0.31, respectively. Most EOPs fell within the expected range of 0.01-1.0, however the ΦCD2301*eBM6 efficiency on strain B32 was at 2.47, which is unusually high, particularly when compared to the WT phage, at 0.0051. Overall, these data demonstrate that such hybrid phage can display some advantages over the parental phage.

Consistent with previous data^14^, the host range of ΦCD2301 and its corresponding exchange mutant ΦCD1801*eBM6, appeared much more limited and each phage was only able to infect 9 panel strains. The propagating strain, CD2301, was exclusively infected by the WT phage and CD7701 was only infected by the mutant phage. As resident prophages can grant immunity to phage infection, PHASTER^40^ analysis was employed to identify possible prophages in the CD2301 genome. A total of seven prophage regions were identified, two of which were scored as intact phages. Despite their presence, no common repressor sequence was identified, suggesting that other host factors may prevent ΦCD1801*eBM6 entry into CD2301, although lysogenic immunity cannot be ruled out. Aside from the propagating strain, the mutant phage was able to infect the remaining 8 strains at an average EOP of 0.148 which was equivalent to the average EOP (0.144) of WT ΦCD2301.

## Discussion

While phage therapy is an attractive alternative to antibiotics for the treatment of CDI, *C. difficile* phages isolated to date have very narrow host ranges, often infecting only a few strains. This compromises their potential application as effective therapeutics. A logical solution would be to use engineering biology to expand the host range. A prerequisite of such an undertaking, and the aim of this study, was to identify and engineer the RBP of existing *C. difficile* phages with a goal of altering their host range.

Prior studies that have examined the interaction between *C. difficile* and phages have focussed on the host receptor^14–18^, leaving the reciprocal phage BRP largely uncharacterised. In phages infecting other species, the tail fibres have been shown ^19–23^ to be the RBP ^19–23^. The experiments undertaken here have confirmed this is also the case for *C. difficile Myoviridae* phages. However, contrary to studies into reprogramming host ranges of other species^41–45^, exchanging the tail fibre alone (*eBM1 mutants) was insufficient to change the host range. Rather, as evidenced from the properties of the *eBM6 mutants, changes to host range were dependent on exchange of the tail fibre together with corresponding downstream hypothetical protein.

The hybrid mutant phages created were able to infect their new target strains, mimicking the host range of the WT phages, except in instances where host interference was suspected. The hypothetical protein, therefore, appears to serve an accessory function to the tail fibre in some capacity and is specific to the fibre of the phage from which it originated. Database searches indicated the encoding gene (*hyp*) to be present in other phages infecting *C. difficile*, but that similarity aside, the encoded polypeptide shares no similarity to any protein of known function. On its own, it is not essential for virion function, although when deleted (ΦCD1801ΔBM5), the number of functional phage particles was severely reduced by almost ∼5000-fold, decreasing the titre by 4-logs of magnitude. Lone exchange of the *hyp* gene had no effect on binding as the ΦCD1801*eBM5 mutant behaved the same as the deletion mutant. The BM5 mutants demonstrated that the hypothetical protein does not determine the host range alone and to function it must be paired with a native tail fibre.

Based on the outcomes of the exchange experiments, a likely function of *hyp* is that of a chaperone-encoding gene. These are typically found immediately downstream of tail fibre genes in other phages ^25,46^, but vary in mode of action and size. Classic chaperones, such as gp38 ^47^ and gp57 ^48,49^ .of T4, and Tfa ^50^ of phage lambda, are typically associated with mediating tail assembly. In contrast, adhesins such as gp38 of T2, K3 and Ox2 phages are displayed at the end of tail fibres and contribute to receptor binding and, therefore, host range ^51,52^. The features observed in the hypothetical protein in ΦCD1801 and ΦCD2301, however, are more consistent with a chaperone that combines both functions. These include the apparent specificity towards its parent tail fibre gene, the lack of tail fibre function in the absence of the hypothetical protein and their large size (496-564aa) compared to typical chaperones (90-200aa). Tfa chaperone-adhesins of phages Mu and P2 have similarly been shown to assist both in tail folding and exhibit specificity to their native tail fibre ^24, 27^.

The host range of any particular phage is dictated by the binding range (determined by phage tail components) and host factors, such as resident prophage elements. The latter can be highly influential but are often and the entire emphasis is placed on the phage components. It is important to consider host factors when working with polylysogenic organisms such as *C. difficile* which can contain up to six prophages, more commonly one to three ^53^. Prophages represent a not inconsiderable obstacle to the successful implementation of phage therapy by preventing infection by therapeutic phages through superinfection exclusion^54^. The current study highlights the advantages of hybrid phages as ΦCD2301*eBM6 was able to overcome this phenomenon in strain CD2315, as its ΦCD2301 backbone isn’t associated with RT078 *C. difficile* strains. Therefore, it is unlikely that these strains contain excluding prophage elements effective against this backbone, for example repressor genes with high sequence similarity. This mutant superseded its WT parent phages by exhibiting the broadest host range, infecting 46.6% of the tested panel and exhibiting the highest EOP of those tested.

The work described here has for the first time shown that the ability of *Myoviridae* phages to bind to their *C. difficile* host is dependent on their tail fibre genes and adjacent *hyp* gene, encoding the RBP and a putative chaperone, respectively. The RBP alone was insufficient for binding but required the presence of the putative chaperone. Capitalising on our discovery and using engineering biology, it proved possible to change the host range of the one phage to that of the other by swapping the respective *tcf/tfp* and *hyp* genes. Significantly, one of the resulting mutants, ΦCD2301*eBM6 surpassed both WT phages with its superior host range and efficiency of infection. This is the first time that genome engineering has been used to successfully expand the host range of a *C. difficile* phage. Our work provides a blueprint for enhancing the host range of *Myoviridae* phages, a prerequisite for implementing phage therapy for the treatment of CDI.

## Online Methods

### Growth of *C. difficile* strains

Required strains of C. difficile (Supplementary, S1) were routinely grown at 37°C in a Don Whitley anaerobic workstation, MG1000 (80% N_2_, 10 CO_2_, 10% H_2_). Overnight cultures were incubated on brain heart infusion media supplemented with 0.5% yeast extract, 0.1% l-cysteine (BHIs); agar plates were additionally supplemented with d-cycloserine (250μg/ml) and cefoxitin (8μg/ml) (*C. difficile* selective supplement, Oxoid, USA; BHIsCC).

### Mutant generation

#### Vector construction

All vectors generated in this study were based on the pRECas1 vector^29^ which contains the RiboCas tool for editing genomes. An aliquot of the purified vector was obtained using a Monarch® Plasmid Miniprep Kit (NEB) from an *E. coli* Top10 (Invitrogen, UK) overnight culture. The vector was the digested using AscI and SalI to linearise it. A homology cassette for each mutant was designed and inserted in its place. The homology cassette consisted of sgRNA and homology arms (+cargo). The sgRNA was designed using Benchling’s CRISPR algorithm (https://benchling.com/) set to *C. difficile* 630 genome, with an NGG PAM targeting genes (*ptp, ptc, tcf, tfp, hyp*) for deletion or sgRN-BM6^55^ for complementation/exchange mutants. The resulting sgRNA was digested with AscI and AatII. The homology arms (HAs) were generated using the primers detailed in Table X and purified CD1801L and CD2301L gDNA as template by PCR of individual components, which were then assembled together by SOEing PCR. The resulting HAs were digested with AatII and SalI. The digested components were ligated and transformed into *E. coli* Top 10, 10-Beta (NEB) or 10-Stable (NEB). The vectors (Supplementary, S2) were screened by PCR, followed by Sanger sequencing, then transferred to conjugation donor strains.

#### Conjugation

Donor strain, *E. coli* sExpress, containing the appropriate vector was grown overnight at 30°C in Luria Bertani (LB) broth, supplemented with 12.5μg/ml chloramphenicol. A 1ml volume of culture was harvested through centrifugation at 1217*g* for 2 min and 30 sec, based on previous approaches^56^. The cells were washed twice in phosphate-buffered saline (PBS). The pellet was transferred into the anaerobic cabinet and mixed with 200μl of overnight *C. difficile*. The mixture was spotted onto BHIs agar and left for 24h. Following that, the growth was collected and resuspended in PBS, then plated on BHIsCC supplemented with 15μg/ml thiamphenicol (BHIsCCTm) to select for transconjugants. The plates were incubated for 72h.

#### Mutant generation and Induction

Following conjugation, the resulting transconjugants were induced to express the RiboCas vector components by plating on BHIsCCTm supplemented with theophylline (48mg/ml), then screened by PCR and Sanger sequencing. Once verified, the strains were continually subcultured to achieve plasmid loss. Plasmid loss was confirmed by screened colonies being unable to grow on BHIsCCTm media, usually after 5 days of culture. Following verification of the mutations again through Sanger sequencing, the mutant *C. difficile* (Supplementary, S3) were induced to produce phages. The *C. difficile* mutants were cultured overnight, then induced with Mitomycin C (3μg/ml) and incubated for 24h. Next day, the cultures were centrifuged at 10000*g* for 10 min to pellet the bacterial cells and debris. The supernatant, containing phages, was sterilised using a 0.22μm filter and stored at 4°C.

### Mutant testing

#### Binding assay

Relevant *C. difficile* strains were subcultured, where a 1% of overnight culture was transferred to 20ml of pre-reduced BHIs broth and incubated for 4h at 37°C, under anaerobic conditions. The 20ml culture was harvested by centrifugation at 10000*g* for 10 min and the pellet was re-suspended in 500μl of BHIs broth. A 100μl aliquot of the resuspended cells was centrifuged again at 3,637*g* for 2 min; the supernatant was discarded, and the cells were resuspended with 100μl of the relevant phage preparation. The phage-cell mixture, alongside phage only controls were incubated for 15 min at 37°C under anaerobic conditions to allow for phage binding to the cells. Following the binding period, the samples were resuspended in 1ml of BHIs broth and centrifuged at 14,549 x *g* for 2 min to pellet the phage-bound cells. The supernatant was sterilised using a 0.22μm filter the resulting sample was enumerated using a standard plaque assay to determine the phage titre in the filtrate. The degree of phage binding was then calculated based on titres (% bound = [[control titre – bound titre] / control titre] x100). The titre is indicative of the number of free phage particles/ml and effectively demonstrates whether phage binding occurs when compared to phage-only control samples.

#### Phage enumeration via plaque assay

Phages were assayed for plaque forming unit (p.f.u.) counts using a double agar overlay plaque assay method^57^. *C. difficile* strains were subcultured, where a 1% of overnight culture was transferred to 20ml of pre-reduced BHIs broth and incubated for 6-8h at 37°C under anaerobic conditions. A 1ml aliquot of this culture was mixed with 200μl of phage (dilution) and 3ml of 0.5% BHIs top agar kept at 50°C. The mixture was poured to form an even layer on BHIsCC base agar plates. Plates were incubated in anaerobic conditions overnight at 37°C and counts were observed the next day, calculating pfu/ml.

#### Transmission Electron Microscopy

Phage particles were prepared using ammonium acetate (Sigma-Aldrich, USA) precipitation as previously established^58^. High titre phage preparation (10^7-9^) was pelleted at 21000*g* for 75 minutes, then washed thrice in 0.1M Ammonium Acetate, the precipitated phage was stored at 4°C. A 13μl aliquot of precipitated phage was placed on a 200-mesh copper grid with carbon/formvar film (EM Resolutions) and stained with 2% uranyl acetate (Sigma-Aldrich, USA) before visualisation under transmission electron microscopy (TEM).

#### Phage host range testing

A standard plaque assay was used to determine the host range of each phage, encompassing a range of dilutions (10^0-6^). Subcultures of each panel strain were incubated with each phage (and dilutions), alongside propagating strains to determine titre, and then calculate the efficiency of plating (EOP). The EOP was calculated by comparing the phage titre obtained using panel strain to the titre of propagating strain (EOP = test strain titre / propagating strain titre).

## Acknowledgments

This work was supported by the Biotechnology and Biological Sciences Research Council (BBSRC), (grant number BB/M008770/1) through an iCASE award with SNIPR BIOME, Copenhagen, Denmark. We thank Denise McLean for TEM assistance.

## Author contributions

J.P.S. and N.P.M. conceived the project; J.P.S. performed the experimental work and analysis; M.J.W. isolated and provided important experimental background of phages ΦCD1801 and ΦCD2301; M.L.K. and S.J.K supported the project; N.P.M. and T.W.B. supervised the research; funding acquisition was by N.P.M.; the manuscript was drafted by J.P.S. with help from S.J.K.; all authors reviewed and revised the paper and agreed to the published version of the paper.

## Competing interests

The authors declare that they have no competing financial interests.

## Additional information

Supplementary information is available for this paper

Correspondence and requests for materials should be addressed to N.P.M.

